# A chemogenetic approach for optical monitoring of voltage in neurons

**DOI:** 10.1101/463109

**Authors:** Mayya Sundukova, Efthymia Prifti, Annalisa Bucci, Kseniia Kirillova, Joana Serrao, Luc Reymond, Miwa Umebayashi, Ruud Hovius, Howard Riezman, Kai Johnsson, Paul A. Heppenstall

## Abstract

Optical monitoring of neuronal voltage using fluorescent indicators is a powerful approach for interrogation of the cellular and molecular logic of the nervous system. Here we describe a Semisynthetic Tethered Voltage Indicator (STeVI1) based upon Nile Red that displays voltage sensitivity when genetically targeted to neuronal membranes. This environmentally sensitive probe allows for wash-free imaging and faithfully detects supra- and subthreshold activity in neurons.

---

Complementation and substitution of electrophysiology methods with non-invasive optical imaging of neuronal activity is a major technological challenge in neuroscience^1,2^. Calcium imaging with genetically encoded indicators is widely used to interrogate the connectivity and function of neural circuits at different spatial and temporal resolutions^2,3^. However, as a surrogate for underlying electrical activity, calcium imaging has a number of shortcomings. For example, calcium indicators lack the sensitivity to register subthreshold activity, and their slow kinetics and the nature of the calcium transient itself often preclude recording of high-frequency firing^2^. Direct readout of neuronal membrane voltage is therefore necessary for imaging subthreshold and inhibitory activity, and for investigating fast-coordinated phenomena^4,5^. As such, significant efforts are being invested in developing probes and improving microscopy for optical monitoring of voltage^6–9^.

Fluorescent indicators for membrane potential can be divided into two groups; synthetic voltage sensitive dyes (VSD)^6^, and genetically encoded voltage indicators (GEVI) based upon voltage sensitive proteins such as opsins, channels and phosphatases^10^. Organic VSDs possess excellent photophysical properties and fast kinetics for live imaging^6,11^. However, their application in vivo is limited by unspecific staining of tissue, compromising signal-to-noise ratio (SNR) and cell identity. GEVIs provide a valuable alternative as they can be genetically targeted to subsets of cells^7,8,12^. However, GEVIs often suffer from low brightness, poor photostability and slow kinetics^7,8^. They may also localize poorly to the plasma membrane and exhibit cellular toxicity. To circumvent these problems, hybrid voltage indicators have been proposed which combine the superior optical properties of small-molecule fluorophores with genetically encoded voltage sensors^13–15^. Examples include a precursor VSD that is converted to an active membrane-bound dye by a genetically encoded enzyme^16^, and click chemistry- and enzyme-mediated ligation of organic fluorophores to rhodopsin to function as FRET donors^17,18^.

Here we considered an alternative approach for hybrid voltage sensor design; localization of a synthetic voltage indicator to cells of interest using genetically encoded protein tags^19,20^. We focused on enzyme-based small protein tags such as the self-modifying enzyme SNAP^21,22^-tag, and transferase-mediated labeling of the acyl carrier protein (ACP)^23,24^-tag, as these technologies allow for rapid, irreversible labeling, and are compatible with in vivo imaging^25,26^. For the VSD component we found that derivatives of Nile Red, an environment-sensitive (‘fluorogenic’) dye that shows fluorescence enhancement upon transition from aqueous to hydrophobic solvent^27,28^, register membrane potential with high fidelity. We named the resulting voltage sensor Semisynthetic Tethered Voltage Indicator 1 (STeVI1).

We initially investigated the voltage sensitivity of the Nile Red-derivative NR12S (Fig.1a), fluorogenic probe which contains a zwitterionic group and hydrocarbon chain, and has been used to monitor lipid order ^29,30^. NR12S readily labeled the membranes of HEK293T cells with a signal-to-noise ratio (SNR) of 10.45±1.3 under wash-free conditions (SI Fig. 1 and SI Fig.4a). To quantify its voltage sensitivity, we used whole cell voltage clamp to control membrane potential, and simultaneously recorded the fluorescence signal from the membranes at an illumination power of 12 mW/mm^2^. Upon application of rectangular depolarizing voltage steps of various magnitudes from a –60 mV holding potential, fluorescence signal decreased linearly in the physiological range of membrane potential (R^2^=0.97, Fig.1b). Voltage sensitivity expressed as fractional fluorescence change ΔF/F% (normalized to fluorescence at a holding potential of – 60 mV) achieved – 5.1 ± 0.4 % per 100 mV (n= 10 cells). NR12S fluorescence responded to applied voltage steps with a mean rise time τ_on_ of 1.9 ± 0.4 ms and a weighted decay time τ_off_ of 1.9 ± 0.2 ms (the fast component represented >85% of response).

**Figure 1.**
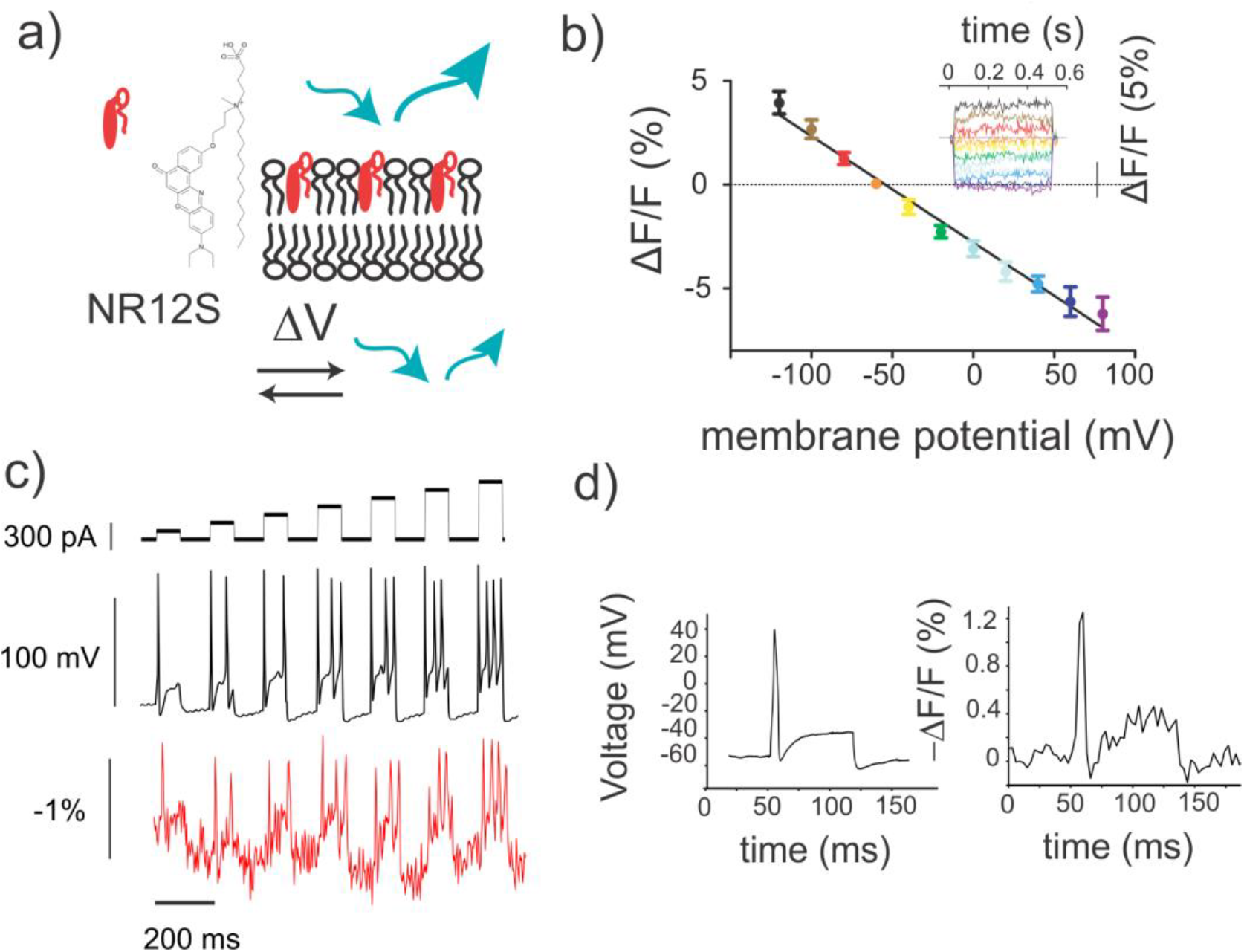
Nile Red based probe NR12S detects membrane voltage change. a) Schematic of Nile Red-based NR12S probe in the membrane. Depolarization of the cell membrane leads to decreased fluorescence intensity. b) Mean response of NR12S to changes in transmembrane potential in patch-clamped HEK293T cells. Error bars represent 95% C.I. (n=6 cells). *Inset graph.* Fluorescence responses in a typical NR12S-labeled HEK293T cell subjected to 500 ms-long square voltage steps of various magnitudes. Responses were normalized to fluorescence at the −60-mV holding potential. Line colors match the colors of data points at the main graph. c) Typical NR12S fluorescence response to single trial recordings of action potentials DRG neurons triggered by current injections of incrementing amplitude (80 ms, 80 - 560 pA, top trace). d) NR12S response to single trial recordings of action potentials in DRG neurons triggered by current injection (80 ms, 600 pA). Full width at half-maximum of the action potential of the voltage trace (left) was 4.5 ms and of its fluorescence readout was 6.3 ms (right). Fluorescence was recorded every 5 ms for b) and 3 ms for c) - d). Light power density was 12 mW/mm^2^. Cells were labeled at 500 nM of NR12S for 7 mins at room temperature.

We further enquired whether NR12S could report action potentials in dissociated dorsal root ganglion (DRG) sensory neurons. Imaging at 333 fps allowed for detection of current injection-triggered action potentials in single trials, with a ΔF/F% per action potential of – 1.9 ± 0.3% (n=4 neurons) corresponding to a peak SNR of 9.5 ± 1.6 (Fig. 1c-d). Because of the large dynamic range of the probe and its fast kinetics, NR12S fluorescence signals closely followed the action potential shape, permitting the optical monitoring of sub- and suprathreshold neuronal events. Moreover, the orange emission of membrane-bound NR12S (λ_max_ = 581 ± 1 nm in live cells, SI Fig. 2), favors this fluorophore over green-emitting VSDs and GEVIs for use in vivo.

We next asked whether Nile Red derivatives could be genetically localized to cells of interest through binding to a protein tag (Fig.2a). We first synthesized Nile Red derivatives that bind to SNAP-tag, and expressed the corresponding tag on the extracellular surface of HEK293T cells via a glycosylphosphatidylinositol (GPI) anchor signal sequence (SI Fig. 3a). Polyethylene glycol (PEG) linkers of n=11 repeats (≈4.8 nm) and charged groups were introduced in the molecules to improve water solubility and reduce nonspecific interactions^31,32^ (SI Scheme 1). Although compounds specifically labeled membranes of cells expressing SNAP-tag, negligible voltage sensitivity was observed as tested by simultaneous patch clamp and imaging (SI Fig. 3c-d). Measurements of emission spectra of the compounds from live cells revealed red-shifted fluorescence in comparison to NR12S (SI Table 1), suggesting that compounds were probing hydrophobic surfaces^33^ of the protein tag rather than the membrane environment.

**Figure 2.**
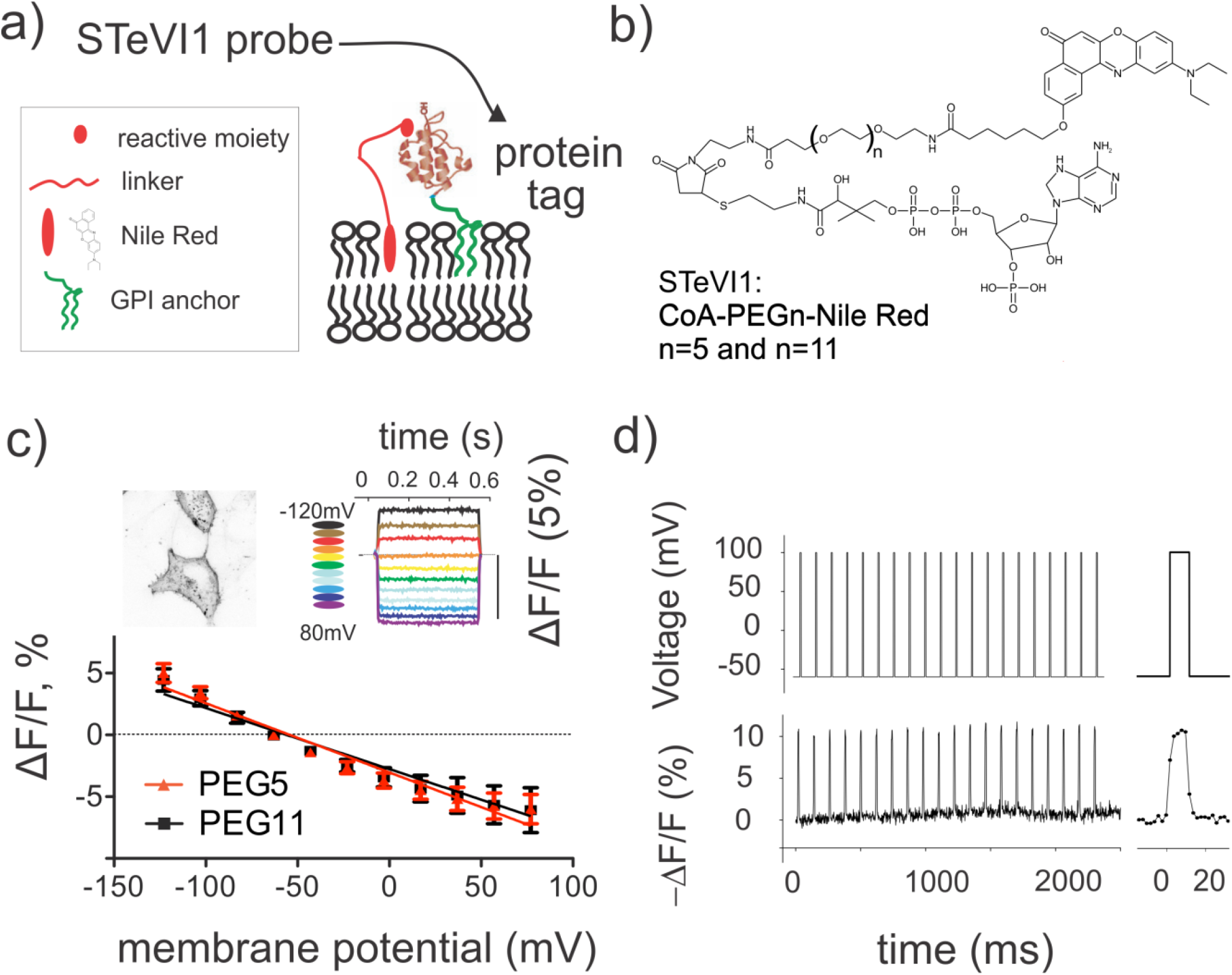
Genetic targeting of Nile Red based voltage indicators. a) Schematic illustration of STeVI1 - Nile Red derivatives tethered with a molecular linker to a protein tag. b) Structures of ACP-targeted Nile Red derivatives with various linker lengths. c) Mean responses of CoA-PEG_n_-NR compounds to changes in transmembrane potential in patch-clamped HEK293T cells, PEG repeats n=5, 11. Error bars represent 95% C.I. (n=8-9 cells). *Inset image.* Representative confocal images of HEK293T cells transfected with ACP-GPI and labeled with 1 μM CoA-PEG_11_-NR. Maximum projection of 19 z-stacks of Δz=0.4μm. LUT is inverted for illustration purposes. Scale bar 10 μm. *Inset graph.* Typical fluorescence responses of CoA-PEG_11_-NR in a HEK293T cell subjected to a 500 ms-long square voltage steps of various magnitudes. Responses were normalized to fluorescence at the −60-mV holding potential. Line colors correspond to different membrane voltages. d) Fluorescence response of CoA-PEG_11_-NR bound to ACP-GPI to rectangular voltage steps of 160 mV, applied at ≈10 Hz (10 ms-duration). Fluorescence recorded every 5 ms for c) and 2 ms for d).

To resolve this issue, we reasoned that a smaller tag may give the probe better access to the membrane electric field. We selected the ACP-tag (8-kDa) and synthesized Nile Red–Coenzyme A (CoA) conjugates with variable PEG repeats (n=5, ≈2.5 nm; n=11, ≈4.8 nm; Fig. 2b, SI Scheme 2). The ACP-tag was expressed on the surface of HEK293T cells via a GPI anchor, and functionality was verified by labeling with CoA-ATTO532 in the presence of phosphopantetheinyl transferase (SFP-synthase, SI Fig. 4d). Nile Red compounds also specifically labeled ACP-GPI expressing HEK293T cells with minimal background (Fig. 2c, SI Fig. 4), SNR of the membrane fluorescence over background under wash-free conditions were 10.7±1.5 for CoA-PEG_11_-NR and 12.4 ±1.3 for CoA-PEG_5_-NR.

ACP-targeted compounds displayed a significant voltage sensitivity in HEK293T cells which, similar for NR12S, was almost linear (R^2^=0.97 and R^2^=0.96 for PEG_11_ and PEG_5_; Fig. 2c and SI Movie 1). Fractional fluorescence change per 100 mV was – 5.5 ± 0.4 % and – 4.9 ± 0.3 % for the CoA-PEG_5_-NR and CoA-PEG_11_-NR compounds (n=9 and n=8 cells). This is comparable to the voltage sensitivity of the non-targetable hemicyanine dye di-4-ANEPPS^34^. We tested the ability of ACP-tag targeted probes to detect trains of action potential-like voltage steps in HEK293T cells (Fig.2d). The kinetics of the CoA-PEG_11_-NR fluorescence response to applied voltage steps was fast, with a mean rise time τ_on_=1.8 ± 0.2 ms and a weighted decay time τ_off_ = 2.6 ± 0.1 ms (the fast component represented >85% of the response).

We next investigated voltage sensitivity of CoA-PEG_n_-NR probes in isolated DRG neurons. ACP-GPI was expressed via recombinant adeno-associated virus (AAV)-mediated gene delivery, and functionality was verified by labeling with CoA-TMR (SI Fig. 5). STeVI1 compounds selectively labeled the neuronal membranes of cell body and axons with negligible intracellular signal (Fig.3a) and no toxicity. Neurons were patch clamped, and action potentials were evoked via current injection. CoA-PEGn-NR compounds tracked single and trains of action potentials in the cell body (Fig.3b,c, SI Fig. 6, SI Movie 2). ΔF/F% amplitudes per action potential reached – 2.0 ± 0.3 % for PEG_11_ and – 2.2 ± 0.3 % for PEG_5_ respectively (n=5 neurons). Corresponding action potential peak SNR were 13.8 ± 1.5 for PEG_11_ and 16.2 ± 3.0 for PEG_5_. ACP-GPI expression and probe insertion in the membrane did not cause significant differences in action potential amplitude and duration, or membrane capacitance between control and labeled neurons (One-Way ANOVA, p=0.11, p=0.72 and p=0.58).

**Figure 3.**
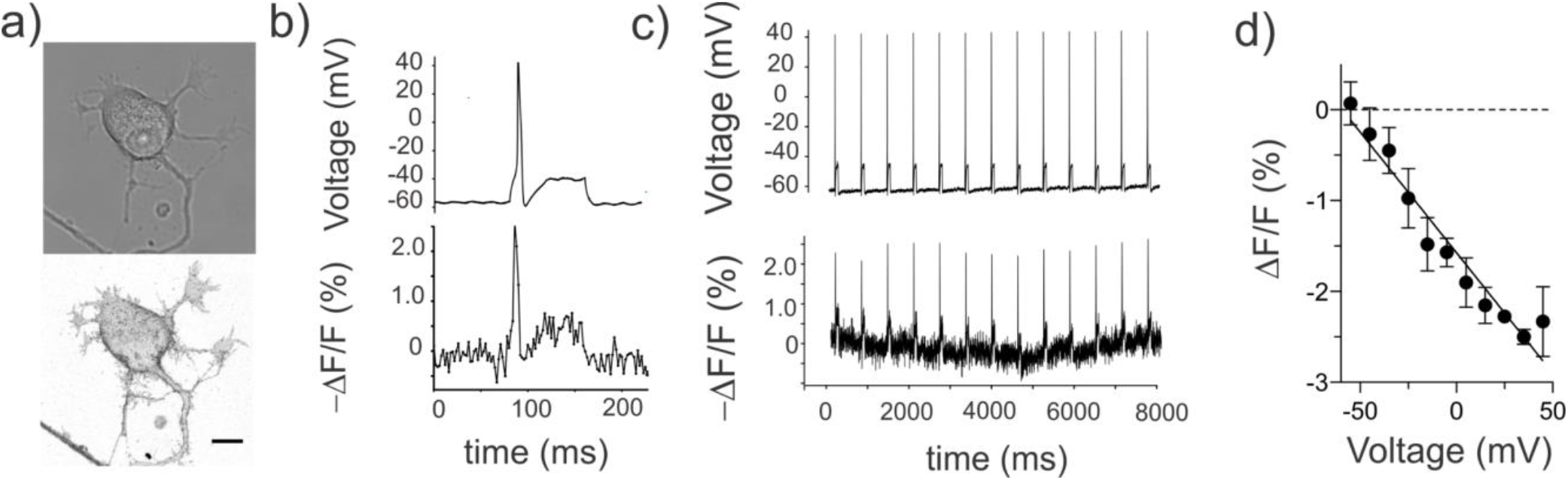
Tethered Nile Red indicators detect evoked and spontaneous neuronal activity. a) Representative confocal images of cultured DRG neurons expressing ACP-GPI via rAAV-mediated delivery. *Top,* brightfield image, *bottom,* maximum projection of 34 z-stacks of Δz=0.4μm. Scale bar 10 μm. b) Typical CoA-PEG_11_-NR fluorescence response in a single trial recording of an action potential in labeled DRG neurons triggered by current injection (80 ms, 80 pA). Full width at half-maximum of the action potential of the voltage trace (top) was 2 ms and of its fluorescence readout was 5 ms (bottom). c) Representative single-trial recordings of current-triggered injection (80 ms, 80 pA) action potentials in DRG neurons with 2 μM CoA-PEG_11_-NR probe bound to ACP-GPI. Fluorescence recorded at 500 fps. d) Fluorescence change vs. membrane potential (mean ± standard deviation) displays the linearity of the voltage sensitivity in neurons (data from the trace in c). Fluorescence changes corresponding to membrane voltage binned to 10 mV intervals were averaged and then fitted with a linear function.

Importantly, in neurons Nile Red fluorescence reported voltage changes linearly (R^2^=0.94, slope = -0.027; Fig. 2d). Fluorescent traces from cell membranes closely mimicked the shape of electrophysiological recorded action potentials, as evidenced by the close match of full width at half-maximum between the fluorescent and electrical traces (Fig. 1c and 3b). Indeed, from the voltage imaging, it was possible to discern the inflection on the falling phase of action potentials that is indicative of nociceptive neurons^35^ (see for example SI Fig. 6). We further investigated whether the Nile Red probes would allow for the detection of spontaneous activity in neuronal processes. Wide-field fluorescence imaging of neurons labeled with CoA-PEG_11_-NR faithfully reported spontaneous spikes at the level of cell body and axons in single-trial optical recordings (Fig. 4, SI Movie 4). ΔF/F amplitude per spontaneous spike at ~10 Hz was -2.3 ± 0.1 % at the cell body membrane corresponding to a peak SNR of 7.6 ± 0.1.

**Figure 4.**
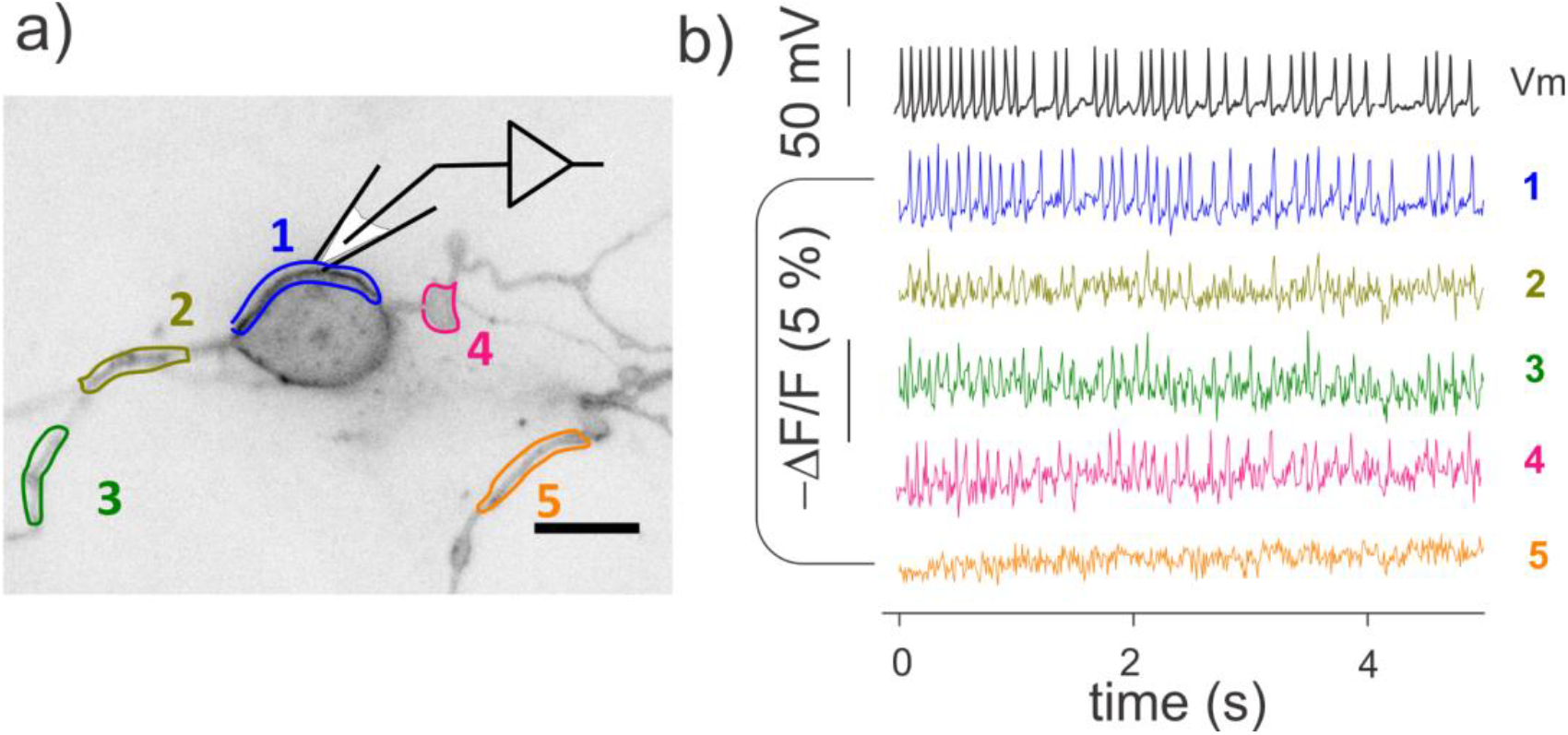
Tethered Nile Red indicators tracks spontaneously occurring action potentials. a) Wide-field image of DRG culture, expressing ACP-GPI and labelled with CoA-PEG_11_-NR. Cell body and individual axons are outlined with colored ROIs. Scale bar is 10 μm. LUT image is inverted. b) Membrane potential (top trace, black) was recorded from the cell body under current clamp mode (no current injection). Fluorescence (measured at 100 fps) from single-trial optical recordings for the color-matched somatic and axon areas (colored traces) highlighted in the image. ΔF/F% reached - 2.3 ± 0.1 % at the cell body membrane. No fluorescence signals were observed for axon area 5 (another neuron).

In conclusion, we demonstrate that Nile Red derivatives display an intrinsic voltage sensitivity that can be exploited to monitor membrane potential in genetically tagged cells. STeVI1 probes were able to detect the shape of subthreshold depolarizations and fast neuronal activity with sensitivity comparable to GEVIs, but required 2-fold less light power (comparison data and references in SI Table 2). Key to the success of the approach was the small size of the ACP-tag which allowed for positioning of the Nile Red in the membrane environment. In future applications, this small size may also enable insertion of the ACP-tag in exposed loops of channels or other membrane proteins, to direct expression to defined subcellular compartments.

## Acknowledgement

We would like to thank EMBL Rome Microscopy Facility, Genetic & Viral Engineering Facility and mouse husbandry Facility. M.S. was supported by Interdisciplinary Postdoctoral Fellowship (EIPOD, EMBL, and EU Marie Curie Actions COFUND II grant) and an award from IBSA Foundation for Scientific Research. E.P. was supported by the European Union Seventh Framework Program (FP7 2007-2013) under grant agreement no. 289278 - “Sphingonet”.

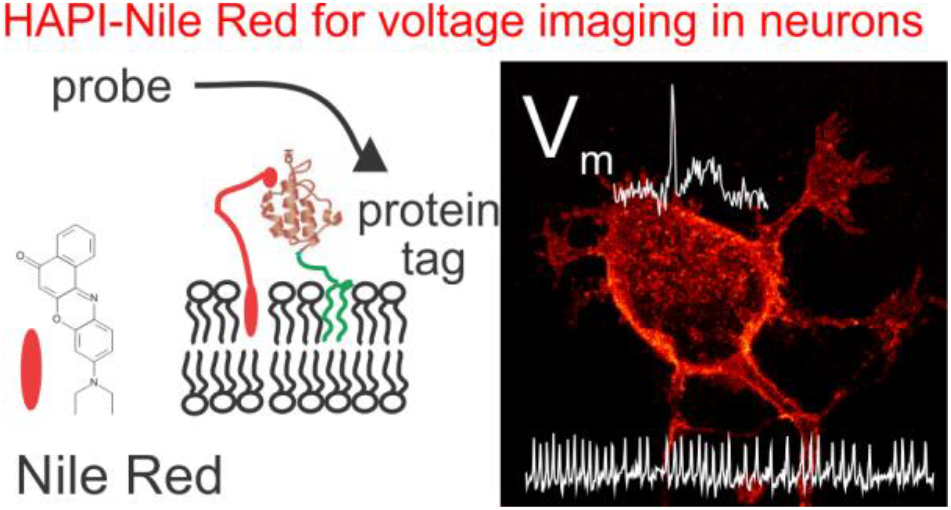

